# Widespread SARS-CoV-2 infection in free-ranging Neotropical bats suggests repeated human-to-bat spillback

**DOI:** 10.64898/2026.08.27.741511

**Authors:** Jorge Armijos-Rivera, Gabriela Parra, Marlon Bravo-Bonilla, María Gonzalez-Maldonado, Daniela Román-Cáceres, Adamary Vásquez-Tituana, Jessica Valdivieso, Freddy Tinoco, Alicia Villavicencio, Nikolay Aguirre, Rodrigo Cisneros-Vidal, Víctor Romero, Jimmy Garces, Domenica de Mora, Alfredo Bruno, Ana Moreno, Clara Tolini, Laura Bavagnoli, Ludvik M. Gomulski, Francesca Scolari, Emmanuele Crespan

## Abstract

Bats harbor exceptional coronavirus diversity and are considered ancestral sources of several human pathogens. As SARS-CoV-2 transitioned from pandemic emergence to global endemicity in humans, concern has shifted from wildlife-to-human spillover toward reverse zoonosis. However, infection of free-ranging bat populations under natural conditions has not previously been demonstrated. Here, we report widespread detection of SARS-CoV-2 RNA in wild Neotropical bats sampled across Andean and Amazonian ecosystems of Southern Ecuador. RT-qPCR screening of 126 individuals, representing nine taxa, detected SARS-CoV-2 RNA in 34.12% of bats across multiple sites. Partial to near-complete viral genomes recovered from five individuals showed >99% nucleotide identity to contemporary human SARS-CoV-2 lineages and clustered within multiple global phylogenetic clades. Mixed-effects modeling revealed pronounced species-level heterogeneity, a positive association between elevation and infection probability, and higher infection probability in females compared with males. The close phylogenetic affinity of bat-derived genomes to circulating human variants and their distribution across multiple lineages suggest repeated anthropogenic spillback rather than sustained bat-specific circulation. These results expand current understanding of the ecological footprint of the COVID-19 pandemic and highlight the importance of integrating wildlife surveillance into long-term One Health strategies for emerging infectious diseases.

## Introduction

Bats are remarkable among mammals for their capacity to harbor a wide diversity of viruses, including SARS-related coronaviruses capable of causing severe disease in humans (El Sayes et al., 2024). They are recognized as important natural reservoirs of numerous zoonotic pathogens, particularly coronaviruses (Letko et al., 2020). Comparative genomic studies indicate that several human coronaviruses - including SARS-related viruses - trace their evolutionary origins to bat-associated lineages, either directly or through intermediate hosts (Wong et al., 2025). These evolutionary patterns, combined with bats’ long lifespans, gregarious roosting behavior, high population densities, global distribution, and their unique immune adaptations (Liu et al., 2022; Yan et al. 2021), create ecological conditions that favor viral maintenance, diversification, and cross-species transmission (Gonzalez & Banerjee, 2022). As a result, bats occupy a central position in discussions of coronavirus emergence and pandemic risk.

The emergence of SARS-CoV-2 in late 2019 and the subsequent COVID-19 pandemic, renewed attention to the evolutionary links between bats and sarbecoviruses (genus Betacoronavirus, family Coronaviridae). Genomic analyses identified closely related coronaviruses in horseshoe bats (genus Rhinolophus), including RaTG13, which shares approximately 96% genome-wide nucleotide identity with SARS-CoV-2, strongly supporting a bat-associated evolutionary origin (Zhou et al., 2020). However, while these findings clarify the evolutionary origins and emergence of the virus, they do not address an equally important and increasingly relevant question: how the virus behaves ecologically after becoming globally established in humans.

As SARS-CoV-2 has become globally endemic in humans, the direction of zoonotic concern has shifted. Beyond initial wildlife-to-human spillover, widespread human infection created unprecedented opportunities for reverse zoonosis, whereby infected individuals transmit the virus back into animal populations. Such spillback events have already been documented in multiple domestic and wild mammals (Milich & Morse, 2024), raising concerns that novel wildlife reservoirs could emerge. The establishment of persistent non-human hosts could enable viral maintenance outside the human population, promote genetic diversification or recombination, and complicate long-term disease control. From both evolutionary and public-health perspectives, the emergence of secondary reservoirs may represent one of the most consequential downstream effects of the pandemic.

Experimental infections have demonstrated that several wild mammals, including Mexican free-tailed bats (Tadarida brasiliensis), are susceptible to SARS-CoV-2 (Porter et al., 2022; Freuling et al., 2020; Bosco-Lauth et al., 2021; Griffin et al., 2021; Hall et al., 2023), indicating that physiological barriers to infection are limited. However, infection of wild bat populations by SARS-CoV-2 under natural conditions has not been demonstrated, as surveillance has largely focused on ancestral bat sarbecoviruses (Pekar et al., 2025) or domestic or peridomestic species (Bashor et al., 2021). Notably, experimental challenge studies in bats have produced species-specific outcomes, ranging from total resistance to transient infection and viral shedding, but with limited evidence for onward transmission to conspecifics (Hall et al., 2023). Therefore, determining whether SARS-CoV-2 can infect natural bat communities is essential, as even short-term infections could facilitate bat-to-bat transmission within dense roosts, expand opportunities for viral evolution across diverse hosts, and promote the establishment of wildlife reservoirs.

Distinguishing susceptibility from reservoir competence is particularly relevant in tropical South America, where exceptional bat diversity coincides with frequent human-wildlife contact. In southern Ecuador, diverse bat assemblages use both natural and human-modified environments, including caves, abandoned buildings, forest edges, agricultural mosaics, livestock areas, and peridomestic structures, which can bring bats into repeated proximity with people and domestic animals, creating plausible opportunities for direct or indirect exposure to human-associated pathogens. Empirical studies have shown that buildings shared by bats and humans generate frequent and prolonged contacts (Jackson et al., 2024), and SARS-CoV-2 risk assessments have specifically identified free-ranging bats as a group of concern for inadvertent human-to-bat transmission (Olival et al., 2020). Although these ecological pathways do not demonstrate transmission by themselves, they provide a strong rationale to test whether human-associated SARS-CoV-2 lineages are present in free-ranging bat communities.

Here, we combine field sampling, molecular diagnostics, genome sequencing, phylogenetics, and ecological modeling to investigate SARS-CoV-2 infection in wild bat communities across Andean and Amazonian ecosystems of Southern Ecuador. We report the detection of SARS-CoV-2 in free-ranging bats and examine ecological and genomic patterns of infection to evaluate alternative transmission scenarios at the human-bat interface.

## Results

### Detection of SARS-CoV-2 in wild bats

In August 2022 and between January and March 2023, we conducted overnight bat sampling at several sites across the provinces of Loja and Zamora Chinchipe in Southern Ecuador, spanning Andean and Amazonian ecosystems (Fig. 1a). In total, 126 bats representing nine taxa were captured, with 20.6% sampled in Loja and 79.4% in Zamora Chinchipe (Supplementary Data 1).

**Fig. 1.**
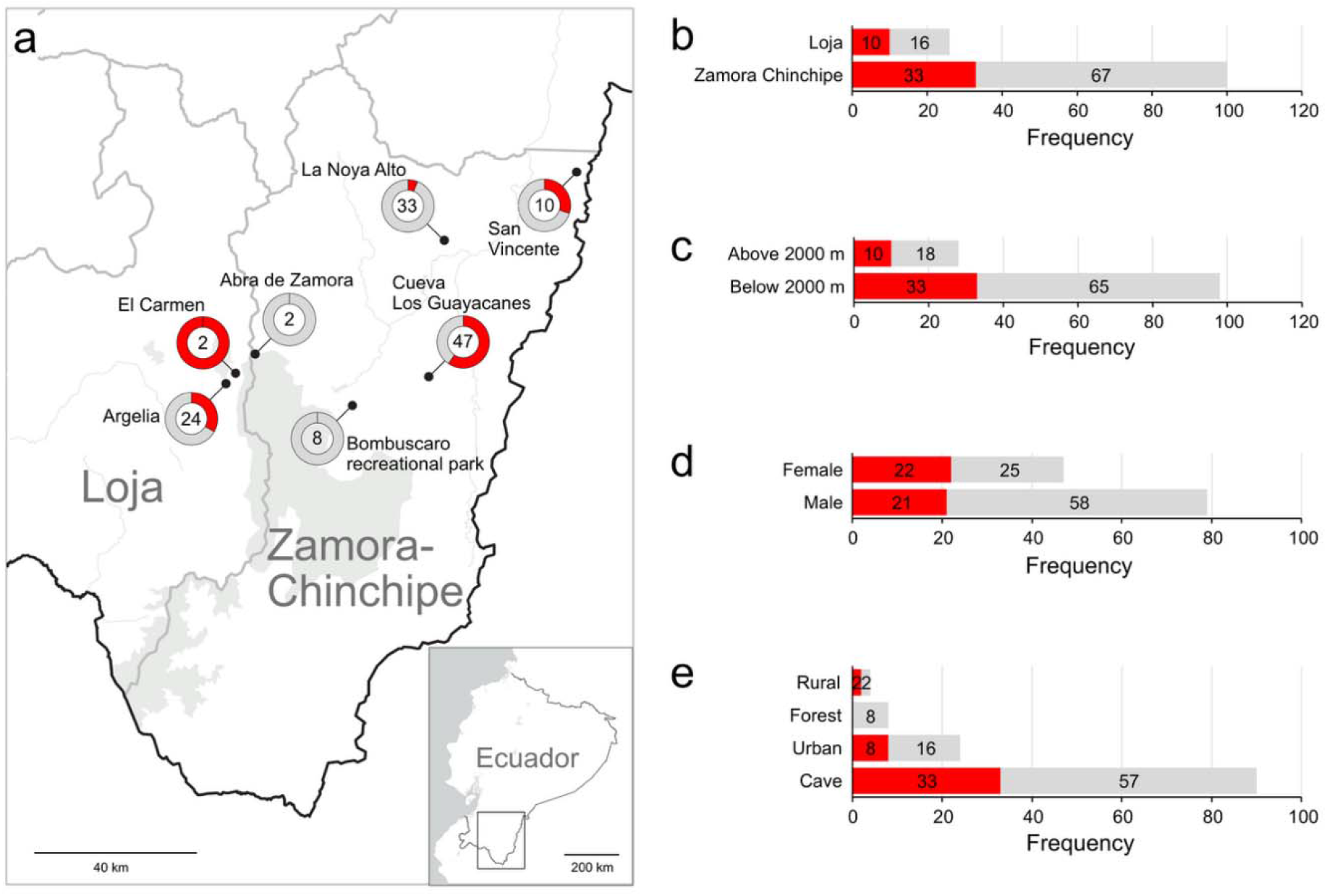
Geographic distribution and ecological correlates of SARS-CoV-2 detection in wild bats from southern Ecuador. (a) Map of bat sampling across Loja and Zamora Chinchipe provinces. Pie charts indicate the proportion of SARS-CoV-2-positive samples (red) and negative samples (grey); sample size is shown at the center of each chart. (b) Proportion of SARS-CoV-2-positive bats by province. (c) Infection prevalence stratified by sample site elevation category (<2000 m vs ≥2000 m above sea level). (d) Infection prevalence by sex. (e) Infection prevalence by habitat type.

Overall, 43 bats (34.12%) tested positive for SARS-CoV-2 RNA by RT-qPCR assays targeting multiple viral genes. The proportion of positive samples appeared to be slightly higher in Loja (38.5%) than in Zamora Chinchipe (33.0%), although this difference was not statistically significant (Fisher’s Exact test, *P* = 0.646; Fig. 1b). Sampling sites spanned elevations from 1,016 to 2,761 m above sea level. When sites were dichotomized at 2,000 m elevation, reflecting well-described environmental transitions in Ecuador (Peters et al., 2010; Zach et al., 2009), infection prevalence did not differ significantly at this coarse elevational scale (Fisher’s Exact test, *P* = 0.825; Fig. 1c).

Sex was associated with infection status: males were significantly less likely to test positive than females (Fisher’s Exact test, *P* = 0.032; Fig. 1d). In contrast, infection prevalence did not differ significantly among habitat categories (Fisher’s Exact test, *P* = 0.1267; Fig. 1e). Positive detections were observed across multiple ecological contexts, supporting the absence of a clear habitat-specific pattern (Fig. 2).

**Fig. 2.**
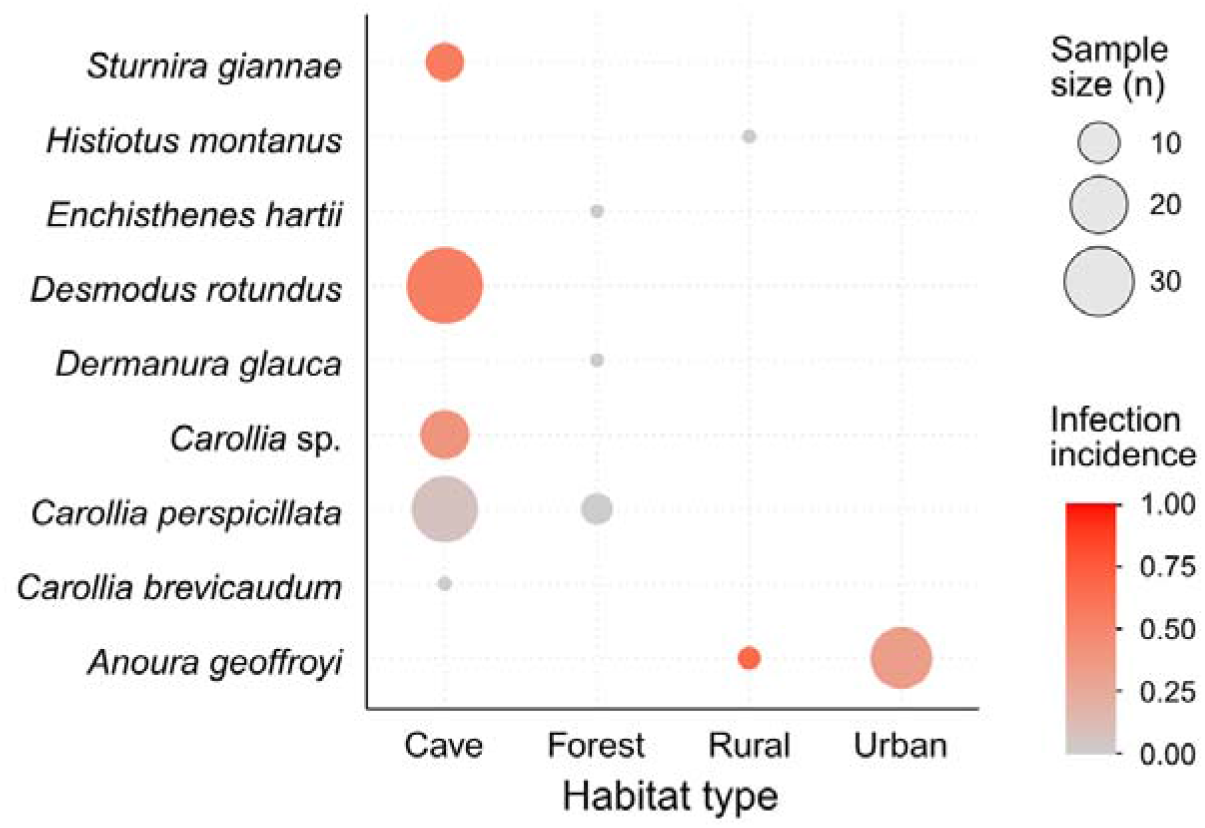
Habitat distribution of bat samples by species and SARS-CoV-2 infection incidence. The circle area is proportional to the sample size. Circle colour indicates infection incidence, ranging from grey (low incidence) to red (high incidence).

Infection probability differed among feeding ecology (binomial Generalized Linear Model, GLM, likelihood ratio test: χ^2^ = 11.9, df = 3, *P* = 0.008). Hematophagous bats showed significantly higher infection odds compared to frugivorous species (odds ratio = 4.34, *P* = 0.001). Nectarivorous bats showed intermediate infection prevalence, although this difference was not statistically significant (*P* = 0.13). Insectivorous bats were excluded from interpretation due to extremely small sample size (n = 1) (Fig. 3). However, in this analysis feeding ecology and species identity were largely confounded, as all hematophagous bats belonged to a single species (*Desmodus rotundus*) and all nectivorous bats were *Ateles geoffroyi*. Therefore, these results should be interpreted as a descriptive contrast among sampled taxon-guild combinations rather than as evidence for an independent effect of feeding guild.

**Fig. 3.**
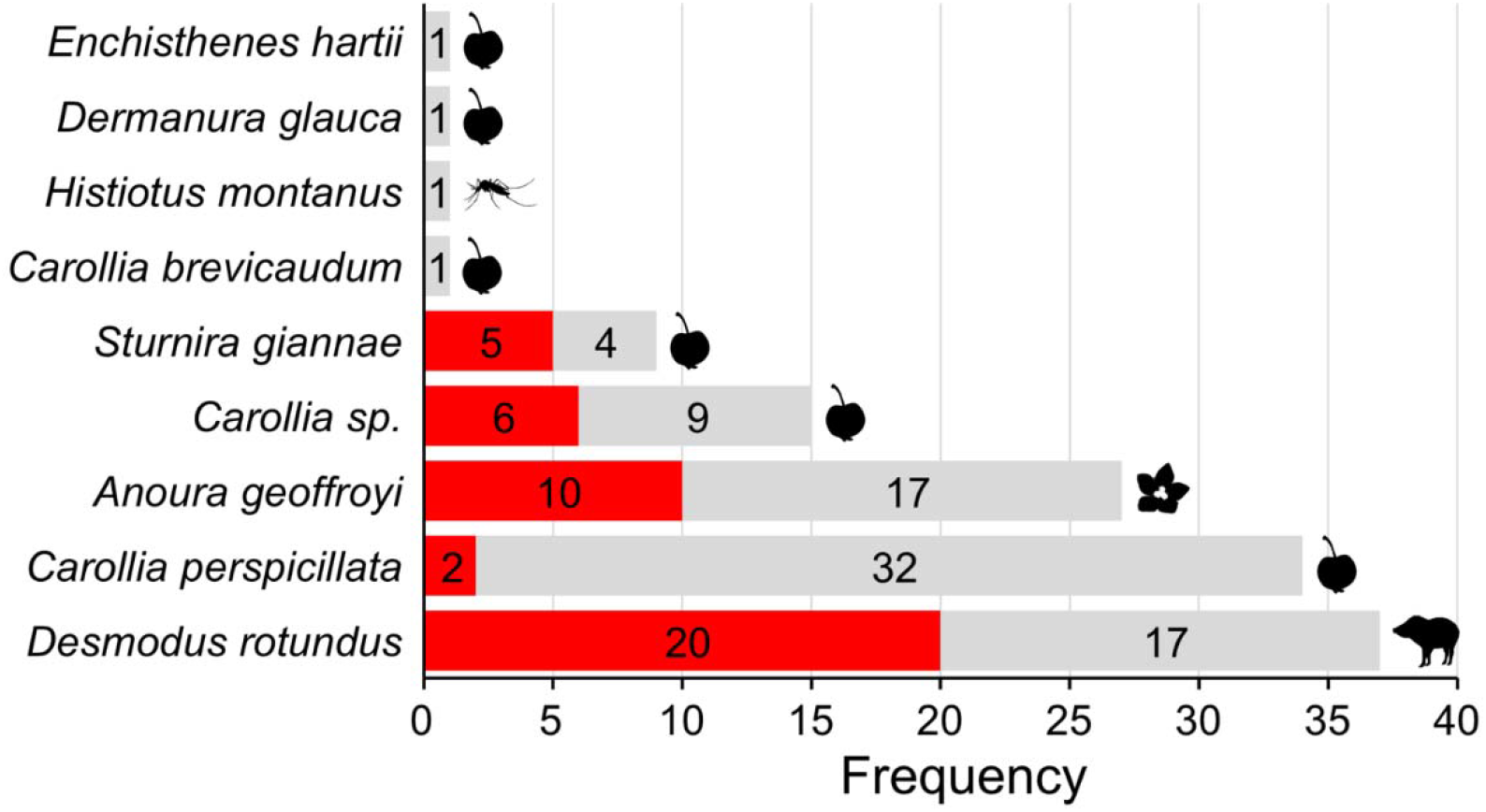
Species-level heterogeneity in SARS-CoV-2 infection prevalence across bat taxa. Observed SARS-CoV-2 infection prevalence by bat species, grouped by feeding ecology. Bars represent numbers of infected (red) and non-infected (grey) individuals. The silhouettes of Malus pumila (CC PDM 1.0), Aedes aegypti by Jaramillo Hector Sergio (CC PDM 1.0), Wahlenbergia by Alexander Schmidt-Lebuhn (CC BY-NC-SA 3.0) and Tayassu pecari by Gabriela Palomo-Munoz (CC BY 4.0) were obtained from PhyloPic (https://www.phylopic.org).

### Species-level heterogeneity and ecological correlations

To account for uneven sampling across taxa and ecological categories, we modeled infection probability using a Bayesian generalized linear mixed-effects framework, including species identity as a random effect (Supplementary Data 2). While the estimated standard deviation of the species-level random effect suggested baseline variation among taxa (posterior mean SD = 0.66; 95% credible interval [CrI]: 0.02-2.23), no individual species differed credibly from the overall mean after accounting for fixed effects, reflecting the high uncertainty driven by uneven sample sizes. Among fixed effects, altitude showed the strongest and most consistent association with infection probability. In the full model - including altitude, sex, feeding ecology, and habitat type - the posterior distribution of the altitude coefficient was clearly shifted away from zero (posterior mean = 1.25 on the log-odds scale; 95% CrI: 0.54-1.97). This corresponds to an approximately 3.5-fold increase in the odds of infection per standard deviation increase in elevation (Supplementary Figure S1). Sex also influenced infection probability after accounting for species identity and environmental covariates: males exhibited lower infection probability than females (posterior mean = −0.94; 95% CrI: −1.82 to −0.09), indicating moderate to strong posterior support for a sex-specific difference in infection risk. In contrast, feeding ecology and habitat type showed limited statistical support as independent predictors of infection probability. Posterior distributions for these effects were centered near zero with wide credible intervals, suggesting that broad ecological classifications did not explain additional variation beyond species identity and altitude.

Given the weak contribution of feeding ecology and habitat, we fitted a more parsimonious multilevel Bayesian model including only altitude and sex as fixed effects, with species retained as a random effect. Results from this simplified model reinforced the primary findings. Infection probability increased strongly with altitude, with an estimated odds ratio of 2.88 per standard deviation increase in altitude (95% CrI: 1.46-6.11). Expressed on the original scale, this corresponds to an approximate 19% increase in infection odds per 100 m increase in elevation. Across the observed altitudinal range, predicted infection probability increased from approximately 10% at the lowest elevations to 85% at the highest elevations (Fig. 4). Importantly, decomposition of altitude into within-species and between-species components demonstrated that the elevation effect was primarily driven by within-species variations rather than differences in species distributions along the gradient. Thus, individuals sampled at higher elevations within the same species exhibited higher infection probability, indicating that the altitude effect cannot be attributed solely to species turnover. Sex differences remained robust in the simplified model. Males exhibited significantly lower infection probability than females (odds ratio = 0.40; 95% CrI: 0.17-0.92), corresponding to predicted infection probabilities of approximately 19% for males and 37% for females at mean altitude (Fig. 4). Species identity was retained as a random effect to account for taxonomic clustering and uneven sampling among taxa. Although the species-level random effect supported this model structure, species-specific deviations should be interpreted cautiously because no species differed credibly from the overall mean after accounting for altitude and sex (all 95% credible intervals overlapped zero on the logit scale) (Fig. 5). This suggests that much of the observed interspecific variation in infection risk may be driven by shared environmental gradients, such as altitude, rather than strong, fixed species-specific differences in susceptibility.

**Fig. 4.**
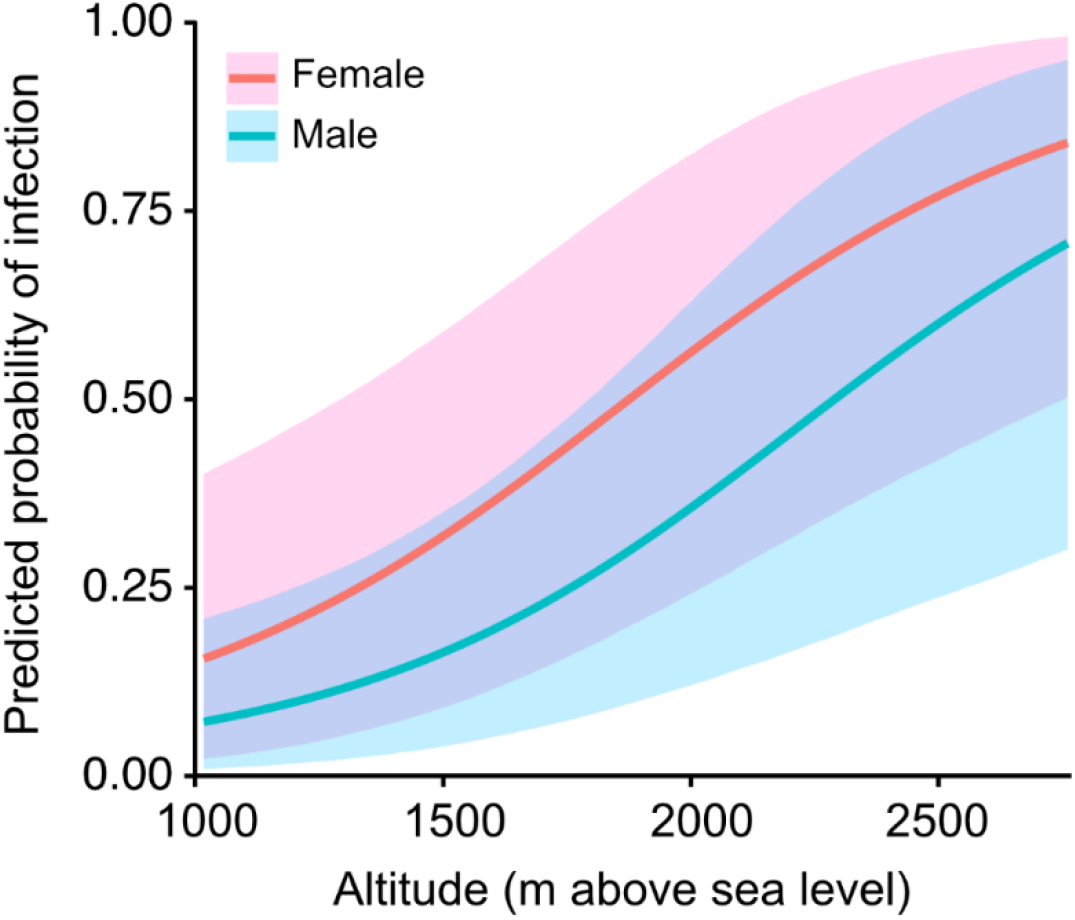
Model-predicted probability of infection as a function of altitude for males and females. Lines show posterior mean predictions from a Bayesian logistic mixed-effects model, and shaded regions indicate 95% credible intervals.

**Fig. 5.**
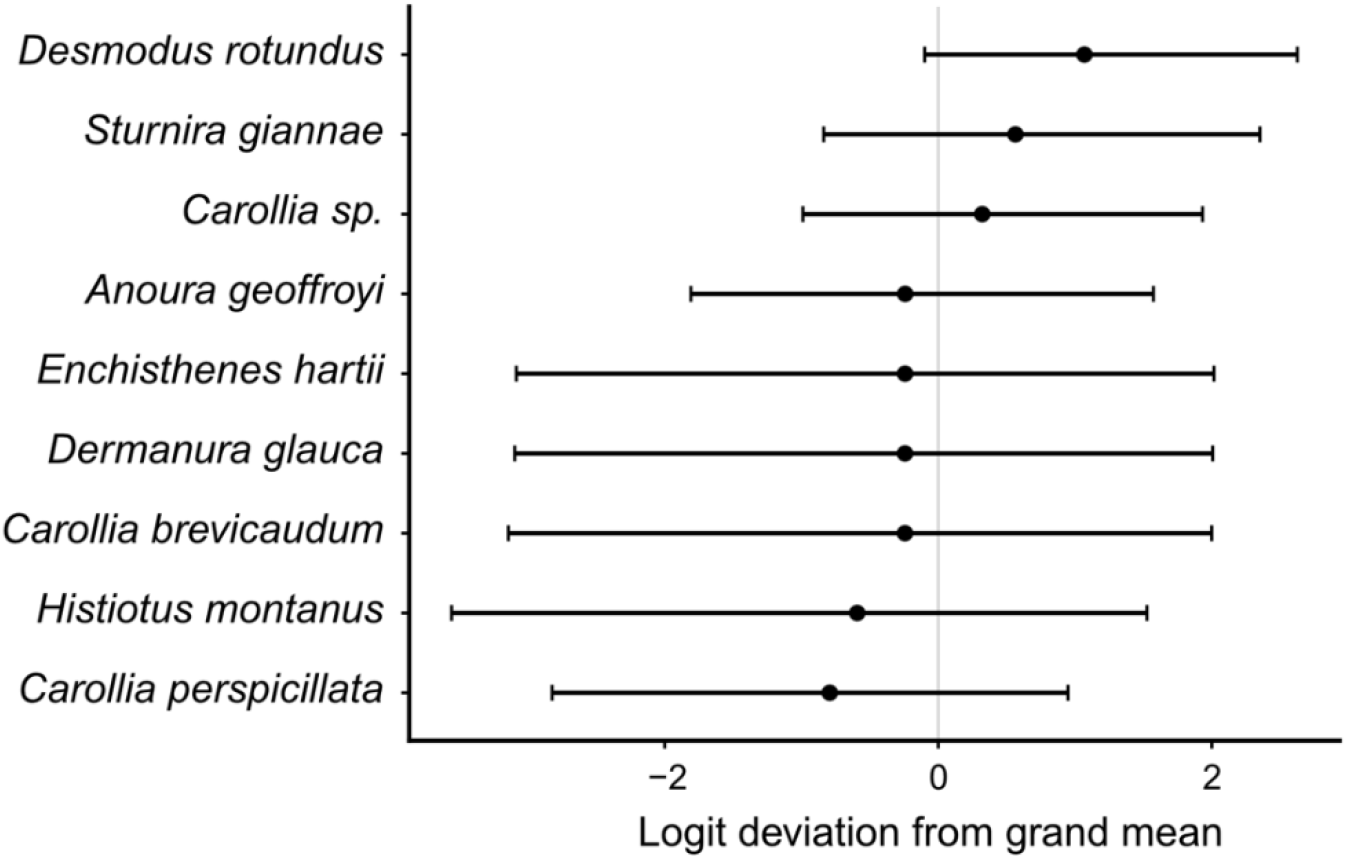
Species-level heterogeneity in infection risk. Posterior means and 95% credible intervals of species-specific random intercepts from a Bayesian logistic model. Values are shown on the logit scale as deviations from the overall mean (0), after accounting for sex and altitude.

### Genomic characterization of bat-derived SARS-CoV-2 sequences

Viral RNA from oral and rectal swabs was subjected to next-generation sequencing. Partial to near-complete SARS-CoV-2 genome sequences were recovered from five positive individuals, including three *D. rotundus* (samples GMB058, GMB060, and GMB068), one *Sturnira giannae* (sample GMB072), and one *Anoura geoffroyi* (sample GMB007). Across these five bat-derived genomes, the number of resolved nucleotide positions ranged from 3,143 to 18,123 after excluding ambiguous sites (Supplementary Data 3). Because genome recovery was uneven among samples, lineage assignments and localized reductions in similarity should be interpreted cautiously considering sequence coverage and the distribution of ambiguous sites.

All five sequences showed high nucleotide identity to the Wuhan genome reference sequence (EPI_ISL_402124), ranging from 99.29% (GMB007) to 99.95% (GMB072) across aligned regions, excluding ambiguous sites and gaps (Stecher et al., 2025). Sliding-window similarity analyses revealed strong conservation across most of the viral genome, with minor localized reductions in similarity within the S and ORF7 regions (Fig. 6). Overall similarity profiles were consistent with limited divergence from globally circulating SARS-CoV-2 lineages.

**Fig. 6.**
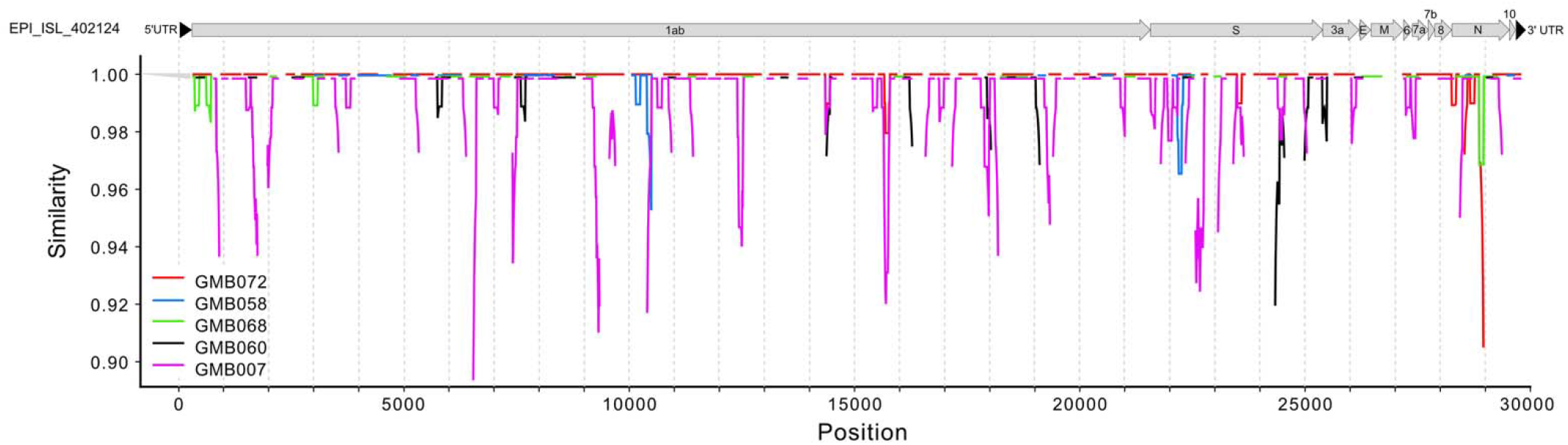
Genome-wide nucleotide similarity between bat-derived and reference SARS-CoV-2 sequences. Nucleotide similarity (%) along the viral genome between the five bat-derived SARS-CoV-2 sequences (GMB072, GMB058, GMB068, GMB060, and GMB007) and the Wuhan reference genome (EPI_ISL_402124), calculated using a sliding window of 100 bp with a step size of 10 bp (Kimura 2-Parameter model). Annotated genomic regions (5′UTR, 1ab, S, 3a, 6, 7a, 7b, E, M, N, 10, and 3′UTR) are shown below the x-axis. Localized reductions in similarity may indicate regions of divergence or recombination among isolates.

### Phylogenetic placement within global and Ecuadorian SARS-CoV-2 diversity

Phylogenetic analyses were conducted using a representative global dataset of 601 SARS-CoV-2 genomes and a second dataset comprising 111 genomes from Ecuador. Maximum-likelihood and Bayesian phylogenies inferred from both datasets showed congruent topologies (Fig. 7, Fig. 8; Supplementary Figures S2 and S3; Supplementary Data 4).

**Fig. 7.**
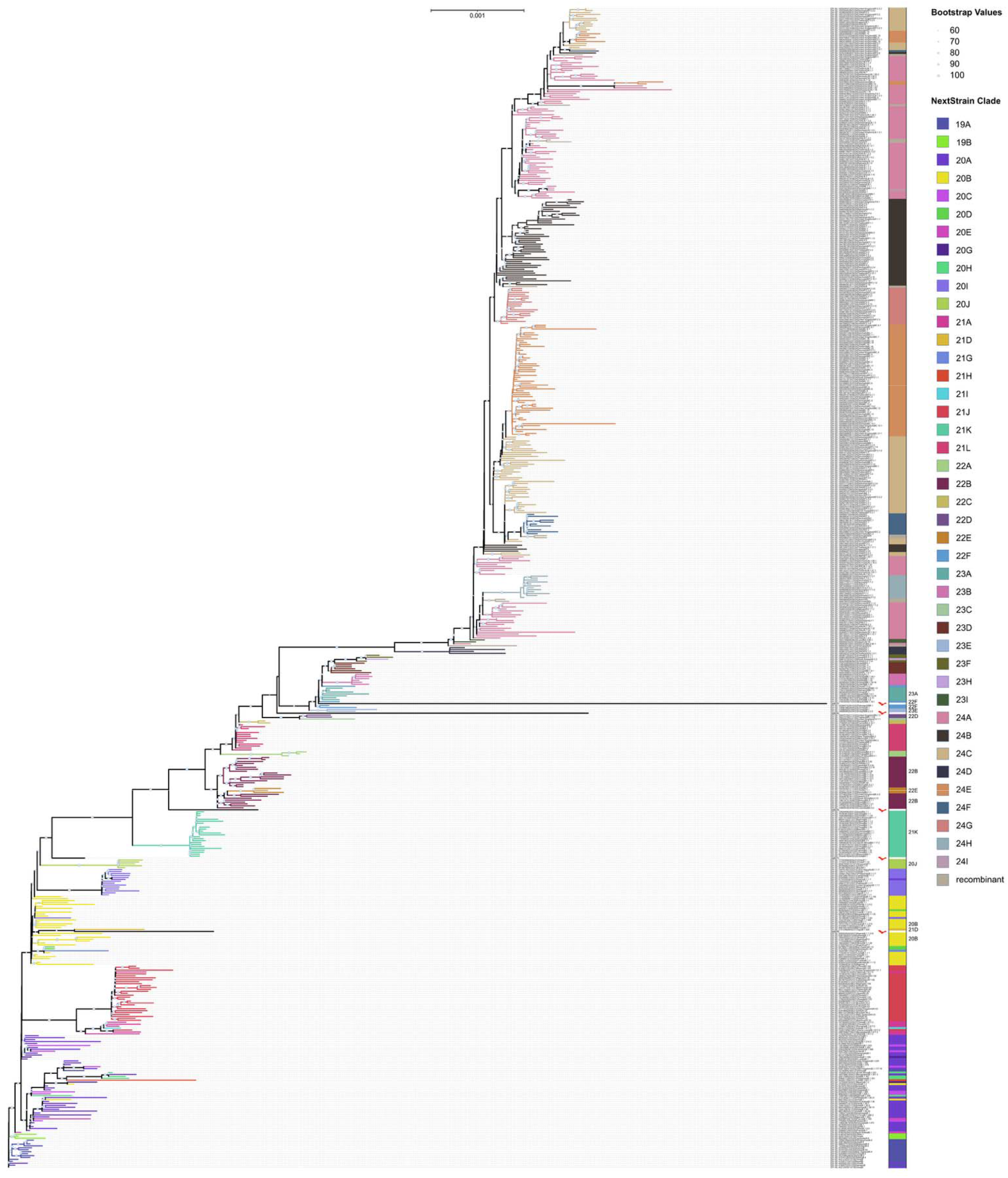
Global phylogenetic placement of bat-derived SARS-CoV-2 sequences. Maximum-likelihood tree of SARS-Cov2 sequences from a representative global dataset of 601 sequences (global). Scale bar indicates the number of base differences per site. NextStrain clades are indicated by colors. The bat silhouette was created by Melissa Ingala (CC BY 3.0) and was obtained from https://www.phylopic.org.

**Fig. 8.**
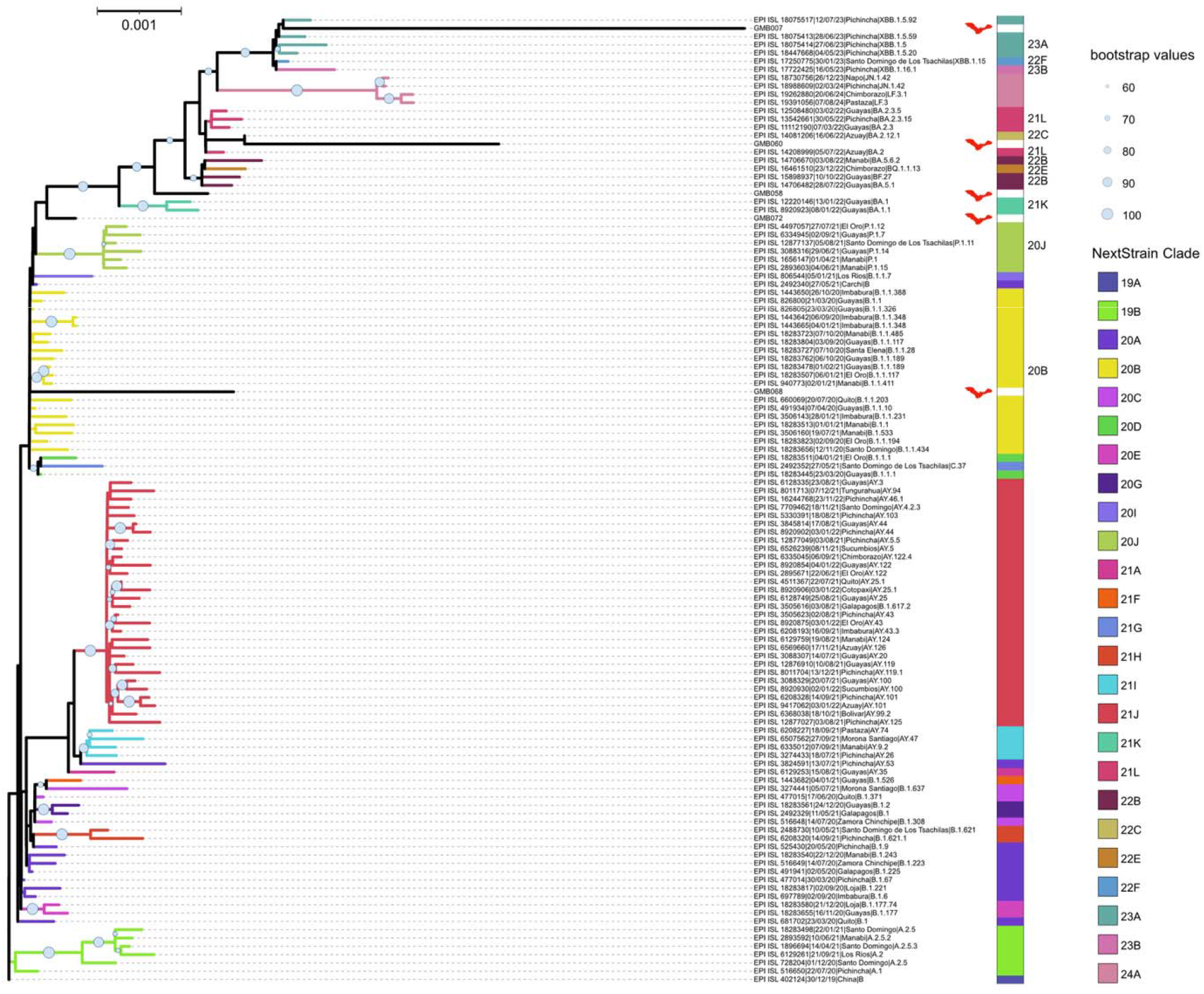
Phylogenetic placement of bat-derived SARS-CoV-2 sequences within Ecuadorian diversity. Maximum-likelihood tree of SARS-Cov2 sequences from Ecuador. Scale bar indicates the number of base differences per site. NextStrain clades are indicated by colors. The bat silhouette was created by Melissa Ingala (CC BY 3.0) and was obtained from https://www.phylopic.org.

In both global and Ecuador-focused analyses, bat-derived sequences clustered within established human SARS-CoV-2 lineages rather than forming a distinct basal clade. Specifically, sequence GMB007 grouped within the XBB.1.x lineage, GMB060 clustered within a BA.2.x clade, and GMB068 fell within the B.1.1.x lineage. The remaining sequences GMB058 and GMB072 occupied intermediate positions between well-supported clades, likely reflecting limited phylogenetic resolution due to partial genome coverage and the reduced number of informative sites. Accordingly, their placement should be interpreted as unresolved or provisional rather than as definitive lineage assignments.

The consistent placement of bat-derived sequences within contemporary human-associated clades across both phylogenetic frameworks indicates close genetic similarity to circulating SARS-CoV-2 variants in Ecuador and globally.

## Discussion

Three major findings emerge from this study: widespread detection of SARS-CoV-2 RNA across multiple bat species, strong phylogenetic affinity between bat-derived and contemporary human lineages, and ecological heterogeneity associated with altitude and sex. In this study, we document widespread detection of SARS-CoV-2 RNA in free-ranging Neotropical bats across multiple species and sites in Southern Ecuador. By integrating molecular screening, genome sequencing, phylogenetics and hierarchical ecological modeling, we show that the viral RNA detected in bats is closely related to contemporary SARS-CoV-2 lineages and is phylogenetically dispersed across distinct clades. Together, these patterns are most consistent with repeated human-to-bat spillback rather than sustained circulation of a bat-adapted lineage.

The scale and duration of SARS-CoV-2 transmission in humans has created unprecedented opportunities for reverse zoonosis, and spillback has now been documented across diverse mammalian systems. In farmed mink, dense populations enabled efficient animal-to-animal transmission and subsequent spillback into humans, demonstrating that animal outbreaks can contribute to onward transmission and potentially generate host-associated genetic signatures (Oude Munnink et al., 2021). In North America, extensive transmission in white-tailed deer has provided compelling evidence that SARS-CoV-2 can persist and evolve outside humans, including cases of accelerated evolutionary dynamics relative to human lineages (Pickering et al., 2022). At a broader community level, recent multi-species wildlife surveillance has shown that SARS-CoV-2 RNA and/or exposure can be detected intermittently across multiple taxa, underscoring that wildlife infections may be more widespread - and more episodic - than initially appreciated (Goldberg et al., 2024). These systems provide an important context for interpreting bat detections: spillback can occur repeatedly, and its consequences range from transient infection episodes to longer-term establishment depending on host ecology and contact structure.

Our phylogenetic results strongly support repeated introductions from humans into bats. The five bat-derived genomes recovered here share >99% identity with human SARS-CoV-2 lineages and cluster within multiple distinct clades rather than forming a single, divergent bat-associated lineage. Such phylogenetic dispersion is difficult to reconcile with one introduction followed by sustained bat-to-bat transmission and progressive divergence; instead, it aligns with a model in which bats are periodically exposed to circulating human variants, producing a mosaic of introductions that reflects the temporal structure of local epidemics. This pattern resembles the “multiple spillovers” framework established for deer, where repeated human-to-animal introductions were followed by onward transmission in some settings (Kuchipudi et al., 2022). In our dataset, however, the absence of a distinct bat clade and the high similarity to contemporary human viruses argue that if bat-to-bat transmission occurs, it is likely short-lived or spatially limited, and does not (yet) produce sustained bat-specific evolution of the virus.

Ecological results further inform plausible transmission scenarios. Infection probability varied strongly among bat taxa, indicating pronounced host-species heterogeneity that may reflect differences in roosting behavior, social structure, immune traits, or exposure pathways. Elevation emerged as the strongest predictor in mixed-effects models even after controlling for species identity, and the within-species elevation effect suggests that this association is not simply explained by different species occurring at different altitudes. Mechanistically, elevational gradients may act as a landscape-level correlate that modulates bat demography, roost density, or microclimatic conditions relevant to viral stability. However, the observed altitude effect may also be partially confounded with unmeasured site-level characteristics that shape encounter rates and exposure opportunities. For instance, lower-elevation sites in this region often coincide with higher human accessibility, livestock interfaces, and activities such as cave tourism, whereas higher or more remote sites feature lower direct human presence and more conserved environments. Therefore, rather than a direct mechanistic driver by itself, elevation likely captures a complex socio-ecological gradient where human movement patterns, settlement connectivity, and local bat assemblage composition intersect to determine transmission risk. Importantly, recent macroecological work predicts that high-elevation and biodiversity hotspot regions may become increasingly important arenas for cross-species viral sharing as climate and land use drive novel species assemblages and contacts (Carlson et al., 2022). Although our study is not designed to disentangle these causal pathways, the elevation signal highlights that reverse zoonosis risk may be structured by landscape context in ways that deserve targeted investigation. These findings highlight the importance of a One Health approach to SARS-CoV-2 surveillance, as increasing interactions among humans, domestic animals, and wildlife may facilitate spillback events and influence their ecological consequences.

Particularly, the pronounced species-level heterogeneity is heavily anchored by the feeding ecology of the common vampire bat (*D. rotundus*), which exhibited significantly higher infection odds compared to frugivorous species. Unlike strictly frugivorous or nectarivorous taxa, the obligate hematophagous nature of *D. rotundus* forces a recurring, direct physical interface with target mammalian hosts, including domestic livestock and humans in anthropized or rural landscapes (Streicker et al., 2016). This unique behavioral bridge significantly elevates exposure opportunities to human-shed pathogens, suggesting this species as an effective sentinel and a primary conduit for anthropogenic spillback across these Neotropical ecosystems.

The observed sex bias, with females showing higher infection probability, is also biologically plausible. In many bat species, females form maternity colonies characterized by dense roosting, prolonged contact, and repeated close interactions during reproductive periods. Such social structure can amplify transmission opportunities following introduction, even if infections are transient. Sex-biased infection dynamics have been shown to scale into population-level effects for wildlife diseases, emphasizing that sex structure can be an important driver of pathogen dynamics and impacts (Kailing et al., 2023).

The detection of SARS-CoV-2-positive bats in cave systems, including caves that are not frequented by humans, requires careful interpretation. At face value, this might seem inconsistent with a spillback hypothesis if one assumes direct and frequent human presence is required at the roost. However, caves may instead function as ecological amplifiers once the virus is introduced elsewhere. Although our data do not demonstrate bat-to-bat transmission, dense roosting, stable microclimates, repeated physical contact, and the co-occurrence of multiple bat species within the same roost provide plausible ecological conditions for within-roost exposure and potential intra- or interspecific spread. This amplification model could allow spillback occurring in more anthropogenic settings, such as near human dwellings, livestock operations, or heavily used landscapes, to later manifest as detectable infections in cave communities without requiring that caves are the primary entry point. Distinguishing such local transmission from repeated independent introductions would require longitudinal roost-level sampling and genomic clustering of viruses recovered from co-roosting individuals.

Our findings should not be interpreted as definitive evidence of a self-sustaining bat reservoir. In particular, the absence of virus isolation, serological evidence, and repeated longitudinal sampling limits our ability to distinguish active infection, recent exposure, and short-lived local transmission. Establishing reservoir status would require evidence of viral persistence independent of human incidence, repeated detection across seasons, and genomic signals consistent with ongoing transmission and host-specific adaptation. The partial nature of several genomes also limits resolution for detecting fine-scale selection or recombination. Nonetheless, even transient infection events may have evolutionary relevance. Recombination is a recognized feature of coronavirus evolution, and SARS-CoV-2 recombinants have been documented in humans, demonstrating that recombination can occur when divergent lineages co-circulate (Jackson et al., 2021).

In principle, periodic spillback into species-rich bat communities—where diverse coronaviruses already circulate—could provide opportunities for ecological and evolutionary interactions involving SARS-CoV-2 over longer timeframes, even if most introductions fail to establish. Crucially, this continuous interaction occurs in an environment where diverse endemic alpha- and betacoronaviruses are known to circulate within Neotropical bat families, multiplying the biological opportunities for heterologous co-infections and subsequent genetic recombination events that could yield novel variants with unpredictable host ranges or phenotypic traits (Wong et al., 2025).

Finally, while contamination can never be completely excluded, several lines of evidence argue against it as a parsimonious explanation: detections occur across multiple sites and taxa; genomes are non-identical; and sequences fall into distinct clades consistent with circulating human diversity. Moreover, the broader literature increasingly supports the plausibility of episodic wildlife infections at the human-animal interface, including in multi-host communities (Goldberg et al., 2024).

## Conclusions

This study provides field evidence that SARS-CoV-2 can be detected in free-ranging Neotropical bats, with phylogenetic patterns most consistent with repeated spillback from humans. Longitudinal sampling and higher-resolution genomic data will be required to determine whether any bat-to-bat transmission occurs and whether it is transient or sustained. As human SARS-CoV-2 circulation persists, reverse zoonosis should remain an explicit component of wildlife disease surveillance, particularly in biodiverse regions where human-wildlife interfaces are frequent. Sustained surveillance that integrates genomic data with fine-scale ecological inference will be essential to distinguish transient spillback from sustained establishment. More broadly, our results suggest that the ecological footprint of the COVID-19 pandemic extends well beyond human populations, potentially reshaping host-virus interactions across diverse wildlife communities.

## Materials and Methods

### Bat sampling and specimen handling

Sampling locations included El Carmen and La Argelia in Loja Canton (Loja Province), as well as San Vicente in Yacuambi Canton, the cave area of Los Guayacanes, La Noya Alto in Yanzatza Canton, and Abra de Zamora and the area near Bombuscaro Recreational Park in Zamora Canton (Zamora Chinchipe Province). At Los Guayacanes and La Noya Alto, captures were carried out near cave entrances identified with the assistance of local residents. Bats were captured using mist nets deployed near cave entrances, abandoned buildings, and forested areas, following established protocols (Kunz & Parsons, 2009). Nets were monitored continuously to minimize handling time and stress. Bats were identified based on external morphological characters using regional taxonomic keys (e.g. Díaz et al., 2021; López Baucells et al., 2016).

### Specimen handling and biosafety procedures

Captured individuals were temporarily placed in thermal bags for a maximum of 10 minutes to allow for species identification. Upon capture, each bat was handled individually and processed separately to minimize stress and prevent cross-contamination between specimens. Individuals were immediately released at the capture site after sampling.

To reduce the risk of human-to-sample contamination and ensure biosafety, personnel wore full protective equipment throughout handling procedures, including disposable coveralls, respirators, and double gloves. Gloves were changed between animals, and handling surfaces were managed to avoid contact between specimens.

### Sample collection and preservation

Oral and rectal swabs were collected from all individuals using sterile calcium alginate swabs according to (Dominguez et al., 2007). Swabs were immediately placed into sterile tubes containing Dulbecco‘s Modified Eagle Medium (DMEM) to preserve RNA integrity and maintain viable virus-infected epithelial cells (Garnett et al., 2020). Each sample was labeled with a unique identifier corresponding to the individual bat and sampling location. Samples were transported on ice from the field to the laboratory and stored at −80°C until RNA extraction. Positive samples were subsequently stored under ultra-freezing conditions prior to downstream sequencing analyses. All sampling and handling procedures complied with Ecuadorian regulations for wildlife research and animal welfare. Field activities and specimen collection were conducted under permits issued by the relevant national authorities, and all protocols followed approved ethical guidelines.

### RNA extraction and molecular screening

All samples were received within 24 hours of collection at a dedicated sample reception area (airlock zone) of the Molecular Biology Laboratory, Centro de Biotecnología, Universidad Nacional de Loja, where they were catalogued and uniquely labelled prior to further processing. Throughout all handling steps, laboratory personnel wore full personal protective equipment (PPE) in accordance with WHO guidelines for the management of potentially infectious biological material, including a full-face respirator fitted with an N95-equivalent particulate filter, double nitrile gloves, a fluid-resistant disposable gown, and a surgical cap (WHO, 2020a). Within the airlock zone, the exterior surface of all transport containers and individual sample tubes was decontaminated with 70% (v/v) ethanol prior to entry into the main laboratory workspace, in accordance with established biosafety protocols for handling potentially infectious specimens (WHO, 2020b). Samples were subsequently placed in secondary containment racks at safe inter-tube spacing to prevent contact between specimens. Tubes were opened individually inside a certified Class II Biological Safety Cabinet (BSC), and each sample was aliquoted into approximately 1 mL volumes in sterile microcentrifuge tubes for long-term storage at −80°C. One aliquot per sample was processed immediately to preserve optimal viral RNA integrity.

Viral RNA was extracted from 200 µL of sample using the MagMAX™ Viral/Pathogen Nucleic Acid Isolation Kit (Applied Biosystems) with automated extraction on an AutoPure Pro32 system (Gentech Biosciences) according to manufacturer instructions. SARS-CoV-2 detection was performed using the TaqPath− COVID-19 CE-IVD RT-PCR Kit (Applied Biosystems) following the manufacturer’s protocol. Briefly, each 20 μl PCR reaction consisted of 6.25 μl TaqPath− 1-Step Multiplex Master Mix 4X, 1.25 μl COVID-19 Real-Time PCR Assay Multiplex, 7.5 μl nuclease-free water, and 10 μl extracted RNA. Amplification targeted the ORF1ab, N, and S genes, with MS2 bacteriophage RNA used as an internal control. Reactions were run on a QuantStudio 5 Real-Time PCR system (Applied Biosystems).

### Resolution of Discrepant SARS-CoV-2 Detection Results

A total of 126 samples were analyzed in this study. Among these, 77 samples were confirmed as negative by both the National Institute of Public Health Research (INSPI; reference laboratory) and the Molecular Biology Lab, Centro de Biotecnología, Universidad Nacional de Loja (CB-UNL; local laboratory). The remaining 49 samples exhibited diagnostic discrepancies between the two institutions and were selected through a predefined and conservative decision framework specifically designed to resolve these inconsistencies. This approach aimed to maximize diagnostic reliability while minimizing the risk of false-positive classification. In the first step, INSPI results were treated as the primary reference. Any sample with confirmed detection of the E gene by INSPI was classified as SARS-CoV-2 positive. For samples reported as negative by INSPI, results obtained at the CB-UNL laboratory were subsequently evaluated using stringent confirmatory criteria. Specifically, samples were classified as positive only when reproducible amplification of at least two independent viral target genes (N and ORF1ab) was observed in two or more technical replicates. Samples showing isolated detection of the S gene or inconsistent amplification across replicates were conservatively classified as negative. This decision reflects the lower diagnostic reliability of single-target detection, the increased susceptibility of the S gene to variability, and the inability to perform additional confirmatory analyses due to sample limitations. Application of this conservative, multi-target decision framework to the 49 discrepant cases resulted in the confirmation of 43 SARS-CoV-2-positive samples and 6 negative samples. Overall, this strategy ensured a high level of confidence in case classification, prioritizing diagnostic specificity and reproducibility in the context of inter-laboratory discrepancies.

### Genome sequencing and similarity analysis

Total RNA was extracted from bat samples using the QIAamp Viral RNA Mini Kit (QIAGEN, Hilden, Germany), according to the manufacturer’s instructions. Amplicon-based next-generation sequencing of SARS-CoV-2 was performed using the Illumina COVIDSeq Assay, 96-sample configuration (Illumina, San Diego, CA, USA), on an Illumina MiSeq sequencing platform, following the manufacturer’s protocol.

First-strand complementary DNA (cDNA) was synthesized from the extracted RNA. SARS-CoV-2 genomic targets were subsequently amplified in two independent multiplex PCR reactions using ARTIC v3 primer pools 1 and 2. The amplicons generated from both primer pools were combined and subjected to tagmentation, adapter and index amplification, purification, and library pooling according to the COVIDSeq workflow.

The final pooled libraries were quantified using the QuantiFluor® ONE dsDNA System on a Quantus− Fluorometer (Promega Corporation, Madison, WI, USA). This fluorescence-based add-and-read assay uses a double-stranded DNA-specific dye with excitation and emission maxima of 504 nm and 531 nm, respectively. Instrument calibration was performed using the supplied Lambda DNA standard, and 1X TE buffer (pH 7.5) was used for standard and sample dilution. Library concentrations obtained by fluorometric quantification were used to normalize the pooled libraries to 4 nM.

The normalized library pool was denatured and diluted to a final loading concentration of 10 pM. Sequencing was performed on the Illumina MiSeq System using a MiSeq Reagent Kit v2 with a 300-cycle configuration and paired-end sequencing of 2 × 150 bp.

Reads were quality-filtered and assembled to generate partial to near-complete SARS-CoV-2 genomes. Sequence similarity to the Wuhan reference genome (EPI_ISL_402124) was assessed using Simplot (Lole et al., 1999) (Simplot++ version 1.3; https://github.com/Stephane-S/Simplot_PlusPlus/releases/tag/v1.3)(Samson et al., 2022)) with a sliding window of 100 bp and a step size of 10 bp.

### Statistical analysis

All statistical analyses were conducted in R version 4.5.1. Infection prevalence in relation to feeding ecology was initially analyzed using a binomial generalized linear model (logit link). The response variable consisted of numbers of infected and uninfected individuals per species, and feeding ecology was included as a categorical predictor. Model significance was evaluated using likelihood ratio tests. Odds ratios were calculated from model coefficients. Subsequent analyses were performed using a Bayesian hierarchical modelling framework. Infection status was treated as a binary response variable (infected/not infected) and analyzed using logistic mixed-effects models implemented in the brms package, which interfaces with Stan via the cmdstanr backend. Individuals with missing infection status or altitude were excluded. Altitude was standardized (mean = 0, SD = 1) prior to analysis to improve model convergence and allow comparison of effect sizes. Sex was included as a fixed effect, and species was included as a random intercept to account for non-independence among individuals and baseline differences in infection probability among species.

Models were fitted using Hamiltonian Monte Carlo with 8 independent chains, each run for 10,000 iterations including 4,000 warm-up iterations. Weakly informative priors were specified for all parameters to regularize estimation while allowing biologically realistic effect sizes. Convergence was assessed using the potential scale reduction factor 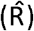, effective sample size, and visual inspection of posterior distributions.

Model fit and predictive performance were evaluated using leave-one-out cross-validation implemented in the loo package, and posterior predictive checks were used to assess agreement between observed and simulated data. To distinguish ecological effects occurring within species from differences among species, altitude was decomposed into within-species (species-centered) and between-species (species mean) components and included simultaneously in an additional model. Posterior distributions were used to estimate effect sizes, credible intervals, and predicted infection probabilities across the observed altitude range, excluding species-level random effects to estimate population-level relationships. Species-specific effects were extracted from posterior random intercept distributions and converted to predicted probabilities by combining random intercept deviations with global intercept.

### Phylogenetic Analysis

A representative global dataset of SARS-CoV-2 genomes was obtained from Nextstrain.org (https://docs.nextstrain.org/projects/ncov/en/latest/reference/remote_inputs.html) on the 17/1/2025. The sequence metadata was filtered to include only sequences with a GISAID EPI_ISL code and good QC score. The dataset was further filtered to include only one example of each Pangolin lineage for each country for each year. This dataset, together with the Wuhan reference sequence (EPI_ISL_402124) and the five bat-derived sequences, was composed of a total of 601 sequences.

A second dataset including only SARS-CoV-2 genomes from Ecuador was assembled using the GISAID repository on 31/1/2025. Only sequences classified as complete, with high coverage and complete collection date were considered. The sequences were filtered to include one example of each Pangolin lineage in each year. This dataset, together with the Wuhan reference sequence (EPI_ISL_402124) and the five bat-derived sequences was composed of a total of 111 sequences.

An acknowledgement for all genomes from GISAID is included in Supplementary Data 4. The sequences were also classified according to the nextstrain clade scheme either directly from the associated metadata or using Nextclade v.3.10.0 (Aksamentov et al., 2021; https://clades.nextstrain.org).

After deleting any ‘N’s in the sequences, they were aligned using MAFFT v7.511 (Katoh et al., 2019) using an iterative refinement method (FFT-NS-i). The resulting alignment was inspected using CLC Main Workbench version 24 (Qiagen Aarhus A/S). Ambiguous bases in the alignment were converted to gaps/missing bases using BBEdit 15.1.3 (Bare Bones Software Inc).

Phylogenetic analyses were performed using Maximum Likelihood (ML; IQ-TREE v. 2.3.6) (Minh et al., 2020) using the model determined within the ModelFinder (Kalyaanamoorthy et al., 2017) option, and Bayesian methods (MrBayes 3.2.7a) (Ronquist et al., 2012) using the model identified by modeltest-ng v. 0.1.7 (Darriba et al., 2020). The trees were visualised using iTOL version 7.0 (Letunic and Bork, 2024) using a palette of optimally distinct colours (https://medialab.github.io/iwanthue/).

The MAFFT alignment of the global SARS-COV-2 dataset was 29,996 nucleotides in length. ModelFinder within IQ-TREE identified the General Time Reversible model (GTR+F+I+R3) (Lanave et al., 1984) as the best fit model and this was used to obtain the optimal Maximum Likelihood tree. A total of 10,000 ultrafast bootstrap replications (Hoang et al., 2018) were performed. For the Bayesian analysis, Modeltest-ng identified the GTR+I+G4 model as being the most appropriate according to both the Bayesian Information Criterion (BIC) and Akaike Information Criterion (AIC), and this choice was confirmed in MrBayes by sampling across the entire general time reversible (GTR) model space in the Bayesian Markov chain Monte Carlo (MCMC) analysis (Huelsenbeck et al., 2004).

The second dataset comprising only SARS-CoV-2 genomes from Ecuador was assembled and aligned, resulting in 29,941 nucleotides in length. The GTR+F+I+R3 model was applied for the ML tree and the GTR+I+G4 model for the Bayesian tree. The ML and Bayesian EcuadorSARS-COV-2 phylogenies (Figure 5 and Supplementary Figure S2) share similar topologies differing only in the placement of some clades near the base of the tree; for example, the sequences belonging to the A.x pangolin clade (19A Nextstrain clade). The positions of the five bat sequences are conserved in the two trees. The GMB007 sequence falls within a XBB.1.x pangolin clade (22F, 23A, 23B Nextstrain clade), GMB060 falls within a BA.2.x clade (21L, 22C), and GMB068 falls within the B.1.1.x (20B) clade. The sequences GMB058 and GMB072 have less well-defined positions. GMB058 falls between a BA.1/BA.1.1 and a poorly supported clade composed of BA.5.1/BA.5.6.2/BF.27 and BQ.1.1.13 sequences (21K and 22B/22E clades), whereas GMB072 falls between the P.1.x and BA.1/BA.1.1 clades (20J and 21K).

### Ethics statement

After identification, all individuals were released unharmed, in strict adherence to animal welfare principles outlined in the Organic Environmental Code (ROS No. 983, Ecuador); the regulations for Zoo-Sanitary Control established by the Agency for Phytosanitary and Zoosanitary Regulation and Control (AGROCALIDAD); and the guidelines for handling wild mammals issued by the Ecuadorian Mammalogy Association (AEM).

## References and Notes

Aksamentov, I., Roemer, C., Hodcroft, E. B., & Neher, R. A. (2021). Nextclade: Clade assignment, mutation calling and quality control for viral genomes. Journal of Open Source Software, 6(67), 3773. 10.21105/joss.03773

Bashor, L., Gagne, R. B., Bosco-Lauth, AM., Bowen, R. A., Stenglein, M., & VandeWoude, S. (2021). SARS-CoV-2 evolution in animals suggests mechanisms for rapid variant selection. Proceedings of the National Academy of Sciences, 118(44), e2105253118. 10.1073/pnas.2105253118

Bosco-Lauth, A. M., Root, J. J., Porter, S. M., Walker, A. E., Guilbert, L., Hawvermale, D., Pepper, A., Maison, RM., Hartwig, A. E., Gordy, P., Bielefeldt-Ohmann, H., & Bowen, R. A. (2021). Peridomestic mammal susceptibility to Severe Acute Respiratory Syndrome Coronavirus 2 infection. Emerging Infectious Diseases, 27(8), 2073–2080. 10.3201/eid2708.210180

Carlson, C. J., Albery, G. F., Merow, C., Trisos, C. H., Zipfel, C. M., Eskew, E. A., Olival, K. J., Ross, N., & Bansal, S. (2022). Climate change increases cross-species viral transmission risk. Nature, 607(7919), 555–562. 10.1038/s41586-022-04788-w

Darriba, D., Posada, D., Kozlov, A. M., Stamatakis, A., Morel, B., & Flouri, T. (2020). ModelTest-NG: A new and scalable tool for the selection of DNA and protein evolutionary models. Molecular Biology and Evolution, 37(1), 291–294. 10.1093/molbev/msz189

Díaz, M. M., Solari, S., Gregorin, R., Aguirre, L. F., & Barquez, R. M. (2021). Clave de identificación de los murciélagos neotropicales. Programa de Conservación de los Murciélagos de Argentina.

Dominguez, S. R., O′Shea, T. J., Oko, L. M., & Holmes, K. V. (2007). Detection of group 1 coronaviruses in bats in North America. Emerging Infectious Diseases, 13(9), 1295–1300. DOI: 10.3201/eid1309.070491.

El Sayes, M., Badra, R., Ali, M. A., El-Shesheny, R., & Kayali, G. (2024). Global Distribution and Molecular Evolution of Bat Coronaviruses. Zoonotic Diseases, 4(2), 146–161. 10.3390/zoonoticdis4020014

Freuling, C. M., Breithaupt, A., Müller, T., Sehl, J., Balkema-Buschmann, A., Rissmann, M., Klein, A., Wylezich, C., Höper, D., Wernike, K., Aebischer, A., Hoffmann, D., Friedrichs, V., Dorhoi, A., Groschup, M. H., Beer, M., & Mettenleiter, T. C. (2020). Susceptibility of raccoon dogs for experimental SARS-CoV-2 infection. Emerging Infectious Diseases, 26(12), 2982–2985. 10.3201/eid2612.203733

Garnett, L., Bello, A., Tran, K. N., Audet, J., Leung, A., Schiffman, Z., Griffin, B. D., Tailor, N., Kobasa, D., & Strong, J. E. (2020). Comparison analysis of different swabs and transport mediums suitable for SARS-CoV-2 testing following shortages. Journal of Virological Methods, 285, 113947. 10.1016/j.jviromet.2020.113947

Goldberg, A. R., Langwig, K. E., Brown, K. L., Marano, J. M., Rai, P., King, K. M., Sharp, A. K., Ceci, A., Kailing, C. D., Kailing, M. J., Briggs, R., Urbano, M. G., Roby, C., Brown, AM., Weger-Lucarelli, J., Finkielstein, C. V., & Hoyt, J. R. (2024). Widespread exposure to SARS-CoV-2 in wildlife communities. Nature Communications, 15(1), 6210. 10.1038/s41467-024-49891-w

Gonzalez, V., & Banerjee, A. (2022). Molecular, ecological, and behavioral drivers of the bat-virus relationship. iScience, 25(8), 104779. 10.1016/j.isci.2022.104779

Griffin, B. D., Chan, M., Tailor, N., Mendoza, E. J., Leung, A., Warner, B. M., Duggan, A. T., Moffat, E., He, S., Garnett, L., Tran, K. N., Banadyga, L., Albietz, A., Tierney, K., Audet, J., Bello, A., Vendramelli, R., Boese, A. S., Fernando, L.,… Kobasa, D. (2021). SARS-CoV-2 infection and transmission in the North American deer mouse. Nature Communications, 12(1), 3612. 10.1038/s41467-021-23848-9

Hall, J. S., Hofmeister, E., Ip, H. S., Nashold, W. S., Leon, A. E., Malavé, C. M., Falendysz, E. A., Rocke, T. E., Carossino, M., Balasuriya, U., & Knowles, S. (2023). Experimental infection of Mexican free-tailed bats (Tadarida brasiliensis) with SARS-CoV-2. mSphere, 8(1), e00263–22. 10.1128/msphere.00263-22

Hoang, D. T., Chernomor, O., von Haeseler, A., Minh, B. Q., & Vinh, L. S. (2018). UFBoot2: Improving the ultrafast bootstrap approximation. Molecular Biology and Evolution, 35(2), 518–522. 10.1093/molbev/msx281

Huelsenbeck, J. P., Larget, B., & Alfaro, M. E. (2004). Bayesian phylogenetic model selection using reversible jump Markov chain Monte Carlo. Molecular Biology and Evolution, 21(6), 1123–1133. DOI: 10.1093/molbev/msh123.

Jackson, B., Boni, M. F., Bull, M. J., Colleran, A., Colquhoun, R. M., Darby, A. C., Haldenby, S., Hill, V., Lucaci, A., McCrone, J. T., Nicholls, S. M., O′Toole, Á., Pacchiarini, N., Poplawski, R., Scher, E., Todd, F., Webster, H. J., Whitehead, M., Wierzbicki, C., … Rambaut, A. (2021). Generation and transmission of interlineage recombinants in the SARS-CoV-2 pandemic. Cell, 184(20), 5179–5188.e8. 10.1016/j.cell.2021.08.014

Jackson, R. T., Lunn, T. J., DeAnglis, I. K., Ogola, J. G., Webala, P. W., & Forbes, K. M. (2024). Frequent and intense human-bat interactions occur in buildings of rural Kenya. PLOS Neglected Tropical Diseases, 18(2), e0011988. 10.1371/journal.pntd.0011988.

Kailing, M. J., Hoyt, J. R., White, J. P., Kaarakka, H. M., Redell, J. A., Leon, A. E., Rocke, T. E., DePue, J. E., Scullon, W. H., Parise, K. L., Foster, J. T., Kilpatrick, A. M., & Langwig, K. E. (2023). Sex-biased infections scale to population impacts for an emerging wildlife disease. Proceedings of the Royal Society B: Biological Sciences, 290(1995), 20230040. 10.1098/rspb.2023.0040

Kalyaanamoorthy, S., Minh, B. Q., Wong, T. K. F., von Haeseler, A., & Jermiin, L. S. (2017). ModelFinder: Fast model selection for accurate phylogenetic estimates. Nature Methods, 14(6), 587–589. 10.1038/nmeth.4285

Katoh, K., Rozewicki, J., & Yamada, K. D. (2019). MAFFT online service: Multiple sequence alignment, interactive sequence choice and visualization. Briefings in Bioinformatics, 20(4), 1160–1166. 10.1093/bib/bbx108

Kuchipudi, S. v., Surendran-Nair, M., Ruden, R. M., Yon, M., Nissly, R. H., Vandegrift, K. J., Nelli, R. K., Li, L., Jayarao, B. M., Maranas, C. D., Levine, N., Willgert, K., Conlan, A. J. K., Olsen, R. J., Davis, J. J., Musser, J. M., Hudson, P. J., & Kapur, V. (2022). Multiple spillovers from humans and onward transmission of SARS-CoV-2 in white-tailed deer. Proceedings of the National Academy of Sciences, 119(6), e2121644119. 10.1073/pnas.2121644119

Kunz, T. H., & Parsons, S. (Eds.). (2009). Ecological and behavioral methods for the study of bats (2nd ed.). Johns Hopkins University Press.

Lanave, C., Preparata, G., Saccone, C., & Serio, G. (1984). A new method for calculating evolutionary substitution rates. Journal of Molecular Evolution, 20(1), 86–93. 10.1007/BF02101990

Letko, M., Seifert, S. N., Olival, K. J., Roche, H., & Munster, V. J. (2020). Bat-borne virus diversity, spillover and emergence. Nature Reviews Microbiology, 18(8), 461–471. 10.1038/s41579-020-0394-z

Letunic, I., & Bork, P. (2024). Interactive Tree of Life (iTOL) v6: Recent updates to the phylogenetic tree display and annotation tool. Nucleic Acids Research, 52(W1), W78–W82. 10.1093/nar/gkae268

Liu X, Li C, Wan Z, Chiu MC, Huang J, Yu Y, Zhu L, Cai JP, Rong L, Song YQ, Chu H, Cai Z, Jiang S, Yuen KY, Zhou. (2022). Analogous comparison unravels heightened antiviral defense and boosted viral infection upon immunosuppression in bat organoids. Signal Transduction and Target Therapy, 7:392. 10.1038/s41392-022-01247-w

Lole, K. S., Bollinger, R. C., Paranjape, R. S., Gadkari, D., Kulkarni, S. S., Novak, N. G., Ingersoll, R., Sheppard, H. W., & Ray, S. C. (1999). Full-length human immunodeficiency virus type 1 genomes from subtype C-infected seroconverters in India, with evidence of intersubtype recombination. Journal of Virology, 73(1), 152–160. 10.1128/JVI.73.1.152-160.1999

López-Baucells, A., Rocha, R., Bobrowiec, P.E.D., Bernard, E., Palmeirim, J.M., and Meyer, C.F.J. (2016). Field guide to Amazonian bats. Instituto Nacional de Pesquisas da Amazônia, Manaus.

Milich, K. M., & Morse, S. S. (2024). The reverse zoonotic potential of SARS-CoV-2. Heliyon, 10(12), e33040. 10.1016/j.heliyon.2024.e33040

Minh, B. Q., Schmidt, H. A., Chernomor, O., Schrempf, D., Woodhams, M. D., von Haeseler, A., & Lanfear, R. (2020). IQ-TREE 2: New models and efficient methods for phylogenetic inference in the genomic era. Molecular Biology and Evolution, 37(5), 1530–1534. 10.1093/molbev/msaa015

Oude Munnink, B. B., Sikkema, R. S., Nieuwenhuijse, D. F., Molenaar, R. J., Munger, E., Molenkamp, R., van der Spek, A., Tolsma, P., Rietveld, A., Brouwer, M., Bouwmeester-Vincken, N., Harders, F., Hakze-van der Honing, R., Wegdam-Blans, M. C. A., Bouwstra, R. J., GeurtsvanKessel, C., van der Eijk, A. A., Velkers, F. C., Smit, L. A. M., … Koopmans, M. P. G. (2021). Transmission of SARS-CoV-2 on mink farms between humans and mink and back to humans. Science, 371(6525), 172–177. 10.1126/science.abe5901

Olival, K. J., Cryan, P. M., Amman, B. R., Baric, R. S., Blehert, D. S., Brook, C. E., et al. (2020). Possibility for reverse zoonotic transmission of SARS-CoV-2 to free-ranging wildlife: A case study of bats. PLOS Pathogens, 16(9), e1008758. 10.1371/journal.ppat.1008758.

Pekar, J. E., Lytras, S., Ghafari, M., Magee, A. F., Parker, E., Wang, Y., Ji, X., Havens, J. L., Katzourakis, A., Vasylyeva, T. I., Suchard, M. A., Hughes, C. C., Hughes, J., Rambaut, A., Robertson, D. L., Dellicour, S., Worobey, M., Wertheim, J. O., & Lemey, P. (2025). The recency and geographical origins of the bat viruses ancestral to SARS-CoV and SARS-CoV-2. Cell, 188(12), 3167–3183.e18. 10.1016/j.cell.2025.03.035

Peters, T., Diertl, K.-H., Gawlik, J., Rankl, M., & Richter, M. (2010). Vascular plant diversity in natural and anthropogenic ecosystems in the Andes of southern Ecuador. Mountain Research and Development, 30(4), 344–356. 10.1659/mrd-journal-d-10-00029.1

Pickering, B., Lung, O., Maguire, F., Kruczkiewicz, P., Kotwa, J. D., Buchanan, T., Gagnier, M., Guthrie, J. L., Jardine, C. M., Marchand-Austin, A., Massé, A., McClinchey, H., Nirmalarajah, K., Aftanas, P., Blais-Savoie, J., Chee, H. Y., Chien, E., Yim, W., Banete, A.,… Bowman, J. (2022). Divergent SARS-CoV-2 variant emerges in white-tailed deer with deer-to-human transmission. Nature Microbiology, 7(12), 2011–2024.10.1038/s41564-022-01268-9

Porter, S. M., Hartwig, A. E., Bielefeldt-Ohmann, H., Bosco-Lauth, A. M., & Root, J. J. (2022). Susceptibility of wild canids to SARS-CoV-2. Emerging Infectious Diseases, 28(9), 1852–1855. 10.3201/eid2809.220223

Ronquist, F., Teslenko, M., van der Mark, P., Ayres, D. L., Darling, A., Höhna, S., Larget, B., Liu, L., Suchard, M. A., & Huelsenbeck, J. P. (2012). MrBayes 3.2: Efficient Bayesian phylogenetic inference and model choice across a large model space. Systematic Biology, 61(3), 539–542. 10.1093/sysbio/sys029

Samson, S., Lord, É., & Makarenkov, V. (2022). SimPlot++: A Python application for representing sequence similarity and detecting recombination. Bioinformatics, 38(11), 3118–3120. 10.1093/bioinformatics/btac287

Stecher, G., Suleski, M., Tao, Q., Tamura, K., & Kumar, S. (2025). MEGA 12.1: Cross-platform release for macOS and Linux operating systems. Journal of Molecular Evolution, 93, 10287. 10.1007/s00239-025-10287-z

Streicker, D. G., Winternitz, J. C., Owen, C. R., Shahi, S., Carter, G., & Gilbert, A. T. (2016). Host-pathogen evolutionary signatures reveal dynamics and future invasions of vampire bat rabies. Proceedings of the National Academy of Sciences, 113(39), 10926–10931. 10.1073/pnas.1606587113

Wong, A. C. P., Lau, S. K. P., & Woo, P. C. Y. (2025). Bats as a mixing vessel for generation of novel coronaviruses: Co-circulation and co-infection of coronaviruses and other viruses. Virology, 604:110426. 10.1016/j.virol.2025.110426

WHO, 2020a. World Health Organization. Rational use of personal protective equipment for coronavirus disease (COVID-19) and considerations during severe shortages. Geneva: WHO. Available at: https://www.who.int/publications/i/item/rational-use-of-personal-protective-equipment-for-coronavirus-disease-(covid-19)-and-considerations-during-severe-shortages

WHO, 2020b. World Health Organization. Laboratory biosafety manual, fourth edition. Geneva: WHO. License: CC BY-NC-SA 3.0 IGO. Available at: https://www.who.int/publications/i/item/9789240011311

Yan H, Jiao H, Liu Q, Zhang Z, Xiong Q, Wang BJ, Wang X, Guo M, Wang LF, Lan K, Chen Y, Zhao H. (2021). ACE2 receptor usage reveals variation in susceptibility to SARS-CoV and SARS-CoV-2 infection among bat species. Nature Ecology and Evolution, (5):600–608. 10.1038/s41559-021-01407-1

Zach, A., Horna, V., Leuschner, C., & Zimmermann, R. (2009). Patterns of wood carbon dioxide efflux across a 2,000-m elevation transect in an Andean moist forest. Oecologia, 162(1), 127–137. 10.1007/s00442-009-1438-2

Zhou, P., Yang, X. L., Wang, X. G., Hu, B., Zhang, L., Zhang, W., Si, H. R., Zhu, Y., Li, B., Huang, C. L., Chen, H. D., Chen, J., Luo, Y., Guo, H., Jiang, R. D., Liu, M. Q., Chen, Y., Shen, X. R., Wang, X., Shi, Z. L. (2020). A pneumonia outbreak associated with a new coronavirus of probable bat origin. Nature, 579(7798), 270–273. 10.1038/s41586-020-2012-7

