## Supplementary Figure S1 for "Widespread SARS-CoV-2 infection in free-ranging Neotropical bats suggests repeated human-to-bat spillback"

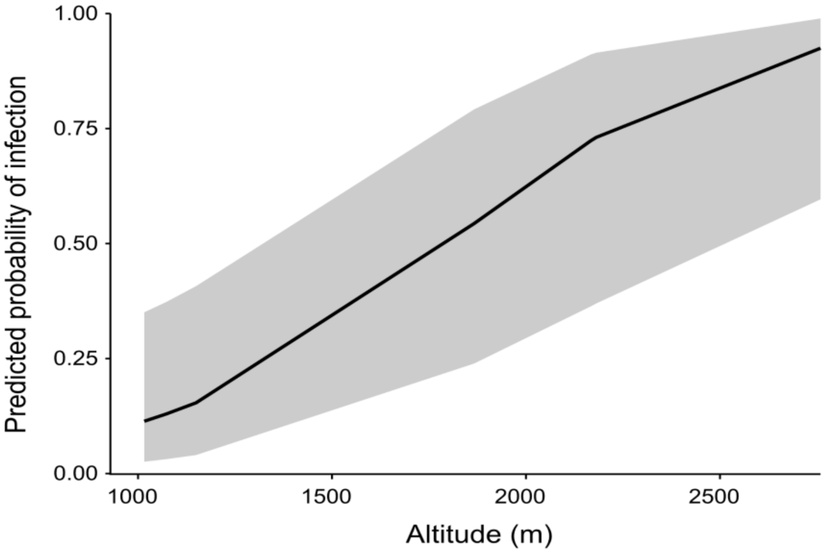


**Supplementary Figure S1**. **Effect of elevation on SARS-CoV-2 infection probability in wild bats.** Posterior estimates from a Bayesian generalized linear mixed-effects model showing the relationship between standardized elevation and infection probability. The line indicates the posterior mean prediction, and shaded region represents the 95% credible interval.
