## Supplementary Figure S2 for "Widespread SARS-CoV-2 infection in free-ranging Neotropical bats suggests repeated human-to-bat spillback"

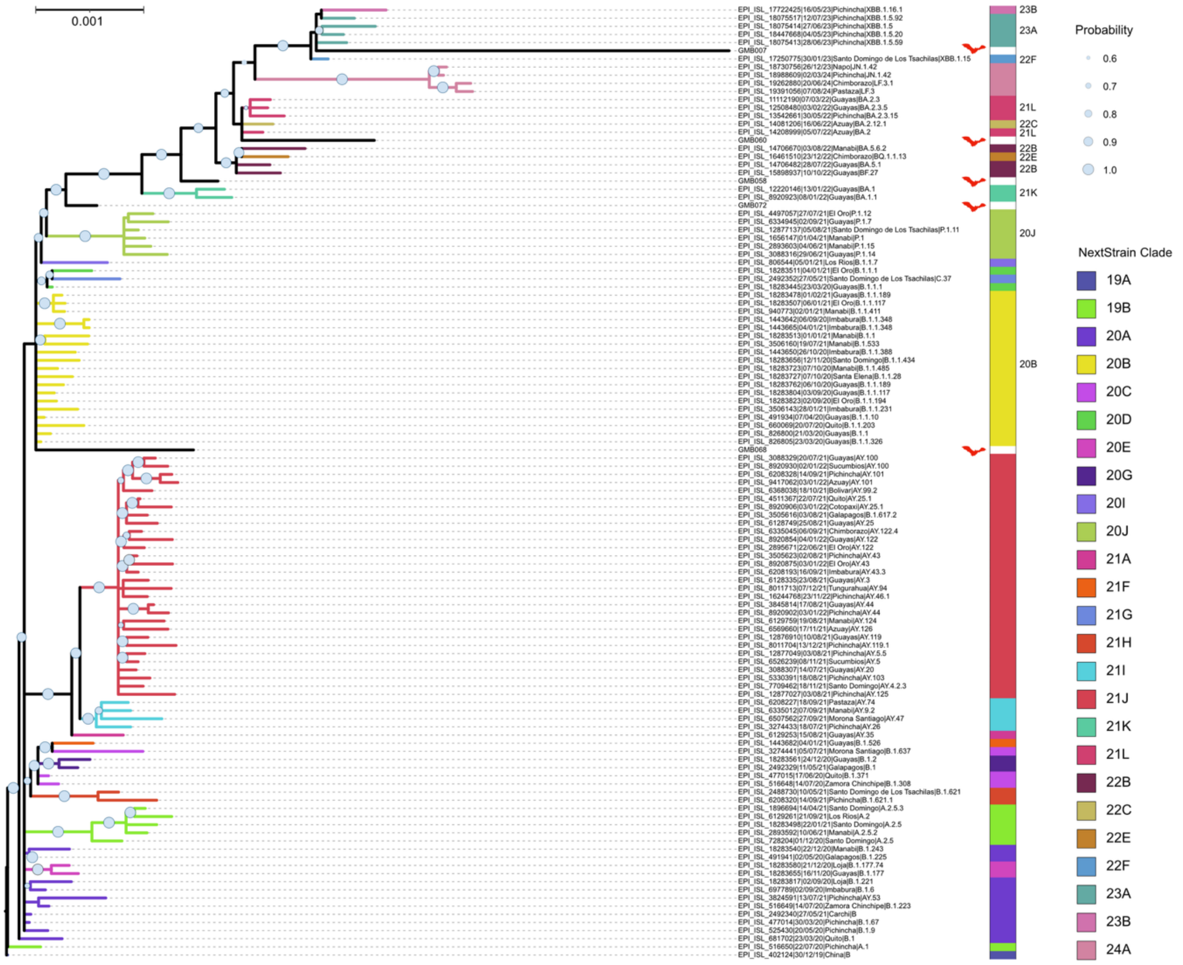


**Supplementary Figure S2.** Bayesian tree of SARS-Cov2 sequences from Ecuador. Scale bar indicates the number of base differences per site. NextStrain clades are indicated by colors. The bat silhouette was created by Melissa Ingala (CC BY 3.0) and was obtained from https://www.phylopic.org.
